# VASH1/2 inhibition accelerates functional recovery of injured nerves

**DOI:** 10.1101/2022.09.20.507919

**Authors:** Philipp Gobrecht, Jeannette Gebel, Alexander Hilla, Günter Gisselmann, Dietmar Fischer

**Affiliations:** Center of Pharmacology, Institute II, Medical Faculty and University of Cologne, Paul-Schallück-Straße 8, 50937, Cologne, Germany; Department of Cell Physiology, Ruhr University of Bochum, Universitätsstraße 150, 44780 Bochum, Germany

## Abstract

Treatments accelerating axon regeneration in the nervous system are still unavailable in the clinic. However, in culture, parthenolide markedly promotes adult sensory neurons’ axon growth by inhibiting microtubule detyrosination. Here, we show that overexpression of vasohibins increases microtubule detyrosination in growth cones and compromises growth in culture and in vivo. Moreover, overexpression of these proteins increases the required parthenolide concentrations to promote axon regeneration, while the knockdown of vasohibins or their enhancer SVBP abolishes parthenolide’s effects, verifying them as pharmacological targets for promoting axon growth. In vivo, repeated intravenous application of parthenolide or its prodrug di-methyl-amino-parthenolide (DMAPT) markedly facilitates regeneration of sensory, motor, and sympathetic axons in injured murine and rat nerves and accelerates functional recovery. Moreover, orally applied DMAPT was similarly effective in promoting nerve regeneration. Thus, pharmacological inhibition of vasohibins facilitates axon regeneration in different species and nerves, making parthenolide and DMAPT promising drugs for curing nerve injury.

## Introduction

Axonal lesions in the peripheral nervous system (PNS) disconnect peripheral targets from the central nervous system (CNS), resulting in motoric, sensory, and autonomic impairments. While PNS axons are, in principle, able to regenerate, functional recovery often remains incomplete because axons can only regenerate with a maximal rate of 1-2 mm/day (1, 2). Moreover, this growth rate can only be maintained for a few months, after which the intrinsic growth potential of neurons is typically lowered, and Schwann cells lose their growth-supporting functions. This time frame limits regenerating axons to overcome maximal distances of approximately 9-18 cm, insufficient for most peripheral nerve injuries (1). Axons that fail to reinnervate their appropriate targets leave patients with permanent functional deficits and cause chronic pain due to inappropriate innervation and misguidance. Therefore, an axon growth rate elevation can quantitatively and qualitatively improve motor and sensory recovery after peripheral nerve injury (3).

Despite extensive research for new treatment strategies, little progress has been made to develop clinically relevant nerve repair drugs. Although factors such as nerve growth factor (NGF) or brain-derived neurotrophic factor (BDNF) accelerate axon growth and neuronal survival in animal models (4, 5), their administration causes severe side effects in humans and therefore is not applicable (Diekmann & Fischer, 2016). Similarly, the immunosuppressor FK506 (Tacrolimus) enhances axonal regeneration upon autologous nerve transplantation (5). However, prolonged and systemic administration of this drug, as required for long-distance nerve injury treatment, inflicts high infection risks, bone fractures, and hypertension (6). Hence, the treatment of peripheral nerve injuries still mainly relies on surgical intervention and depends on the severity of the damage (5, 7). While nerve contusions are usually not treated and, to some extent, heal spontaneously, fully severed nerves normally are re-adapted at both ends. Nerve gaps are often bridged with autologous nerve transplants, which require sacrificing healthy nerves. The sural or the antebrachial cutaneous nerve, typically utilized as transplants, leads to numbness of the respective outer foot or inner arm. Lately, synthetic nerve guides can be implanted into short lesion sites, but so far, these enable insufficient nerve regeneration (5). Surgical interventions to re-adapt severed nerves cannot sufficiently solve slow and often incomplete functional recovery.

Increased glycogen synthase kinase 3 (GSK3) activity in genetically modified mice markedly raises peripheral nerves’ axon growth rate. Furthermore, it accelerates functional recovery, likely mediated by maintaining microtubules in a dynamically unstable state (8). This effect is conveyed by microtubule-associated protein 1B (MAP1B) phosphorylation, which reduces tubulin detyrosination through interaction with tubulin tyrosine ligase (9). In addition, MAP1B has been shown to prevent microtubule detyrosination by direct binding to tyrosinated tubulin (10). Therefore, it is assumed that active GSK3 reduces tubulin detyrosination by phosphorylating MAP1B, thereby increasing microtubule dynamics and, consequently, axonal growth (8, 10–13).

Since genetic alterations are not yet suitable for clinical application, we tested whether pharmacological approaches can similarly reduce microtubule detyrosination in peripheral growth cones and identified the sesquiterpene lactone parthenolide as a molecule able to induce such effects on microtubules and promote the axon growth of adult sensory neurons in culture (11). However, the mechanism underlying parthenolide’s effect on microtubule detyrosination and axon growth remained elusive because the identity of the mediating carboxypeptidase(s) responsible for detyrosination was unknown. However, recently, vasohibin 1 and vasohibin 2 (VASH1, VASH2) were identified as the first two tubulin carboxypeptidases (14–16). Although parthenolide modulates their activity (16), the extent to which VASH1 and VASH2 are involved in the detyrosination of tubulin during axon regeneration is not yet known. This is particularly true because at high concentrations, parthenolide reportedly also forms adducts on both cysteine and histidine residues on tubulin itself, suggesting an indirect inhibition by reducing the polymerization-competent pool of tubulin (17).

The current study provides evidence that vasohibins are the carboxypeptidase involved in axon regeneration and the molecular targets of parthenolide in this context. Moreover, the systemic application of parthenolide or DMAPT accelerates functional motor and sensory recovery in different nerves and species. Thus, inhibition of vasohibins by parthenolide or DMAPT is a promising and potentially clinically feasible approach to treating nerve injuries.

## Results

### Neuronal overexpression of VASH1/2 induces microtubule detyrosination and compromises axon growth of primary neurons

VASH1 and VASH2 were identified as potential carboxypeptidases interacting with parthenolide (15–17). However, their role as molecular targets in promoting axon regeneration remained elusive. To test whether VASH1 and VASH2 play a role in this context, we virally overexpressed both enzymes or their common enhancer SVBP in adult sensory neurons. Then we measured the effects on microtubule detyrosination in axonal tips and, functionally, on axonal growth. As described previously (18), expression of the HA- or FLAG-tagged proteins was detected in neurons 12-24 h after baculoviral transduction. Furthermore, overexpression of either VASH1-HA or VASH2-HA significantly increased microtubule detyrosination in axon tips after 48 h in culture compared to GFP-transduced controls (Fig. 1 A, B), indicating their activity as tubulin carboxypeptidases in neurons. In contrast, SVBP overexpression alone showed no significant effect (Fig. 1 A, B), indicating that additional protein does not enhance detyrosination further. Consistently, the overexpression of each VASH alone, but not SVBP, reduced the average length of regenerated axons (Fig. 1 C, D).

**Figure 1:**
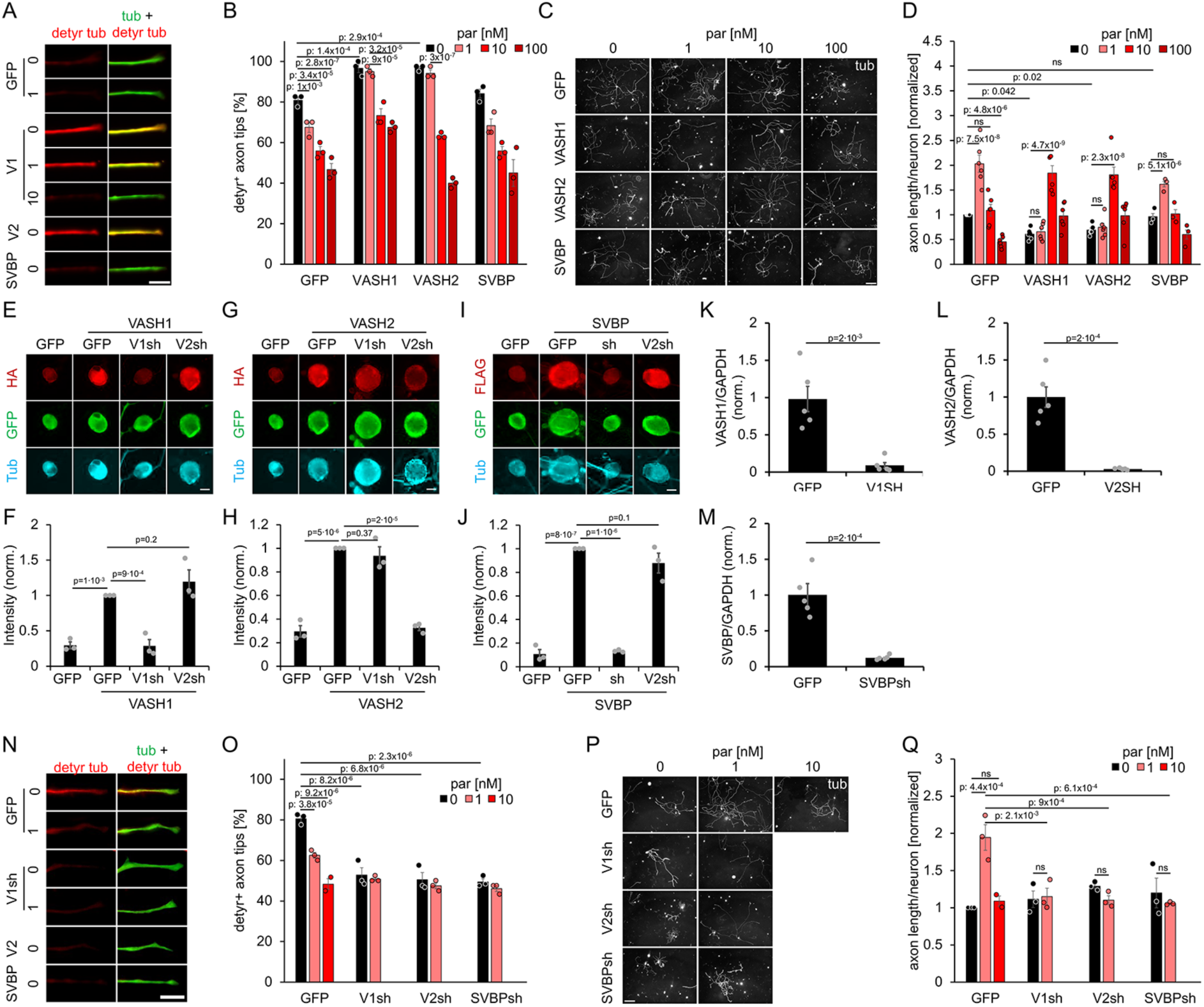
Parthenolide mediated VASH 1, and VASH inhibition promotes axon growth. **A)** Representative photos of axon tips of adult mouse sensory neurons after 2 days in culture, baculovirally transduced with GFP, VASH1 (V1), VASH2 (V2), or SVBP and treated either with 0, 1, 10 or 100 nM of parthenolide. Axons were stained for detyrosinated tubulin (detyr tub, red) and βIII-tubulin (tub, green). Scale bar, 5 μm. **B)** Quantification of detyrosinated tubulin in axon tips of cultures described in A exposed to 0, 1, 10, or 100 nM of parthenolide. Data represent the mean ± SEM of three independent experiments. P-values were determined using two-way ANOVA followed by the Holm-Sidak post hoc test. **C)** βIII-tubulin-immunostained neurons, as described in A. Scale bar, 100 μm. **D)** Quantification of axon growth as depicted in C. Data were normalized to vehicle controls with an average axon length of 543 μm/neuron. Data represent the mean ± SEM of three independent experiments. P-values were determined using two-way ANOVA followed by the Holm-Sidak post hoc test. **E, G, I)** Representative photos of immunostained mouse sensory neurons after baculoviral transduction with GFP, VASH1-HA, VASH-2-HA, or FLAG-SVBP and 2 days in culture. Cultures were co-transduced with either VASH1 shRNA (V1sh), VASH2 shRNA (V2sh), or SVBP shRNA (sh). Neurons were stained for HA or FLAG (red), GFP (green), and βIII-tubulin (tub, cyan). Scale bar, 20 μm. **F, H, J)** Quantification of HA- or FLAG-immunostaining intensity in sensory neurons, as described in E, G, I. Data represent the mean ± SEM of three independent experiments. P-values were determined using two-way ANOVA followed by the Holm-Sidak post hoc test. **K, L, M)** Verification of baculoviral induced knockdown of K) VASH1, M) VASH2, and N) SVBP in quantitative real-time PCR using cDNA from cultivated sensory neurons. For this experiment, sensory neurons isolated from six animals were pooled. All samples were measured five times. Data represent the mean ± SEM of three independent experiments. P-values were determined using two-way ANOVA followed by the Holm-Sidak post hoc test. **N)** Representative photos of axon tips of mouse sensory neurons virally transduced with either GFP or shRNAs against VASH1 (V1sh), VASH2 (V2sh), or SVBP (SVBPsh). Neurons were replated after 5 days and treated with 0 or 1 nM of parthenolide. Regenerated axons were stained for detyrosinated tubulin (detyr tub, red) and βIII-tubulin (tub, green) after an additional 24 h in culture. Scale bar, 5 μm. **O)** Quantification of detyrosinated tubulin in axon tips of sensory neuron cultures, as described in N. Data represents the mean ± SEM of three independent experiments. P-values were determined using two-way ANOVA followed by the Holm-Sidak post hoc test. **P)** βIII-tubulin-stained neurons from cultures described in N were treated with either 0, 1, or 10 nM of parthenolide. Scale bar, 100 μm. **Q)** Quantification of axon growth of neurons as depicted in P. Data were normalized to vehicle controls with an average axon length of 136 μm/neuron. Data represent the mean ± SEM of three independent experiments. P-values were determined using two-way ANOVA followed by the Holm-Sidak post hoc test.

We then tested whether these VASH1/2 or SVBP overexpressing neurons were still responsive to parthenolide. Neurons transduced with a GFP-control virus showed the most robust parthenolide-induced axon growth at 1 nM. At the same time, higher concentrations were less effective, confirming the previously described bell-shaped concentration-response curve (Fig. 1 C, D) (11). Consistently, a parthenolide concentration of 100 nM reduced axon growth due to a more pronounced and prevailing microtubule destabilization at high concentrations (Fig. 1 C, D). Thus, a defined equilibrium range of tyrosinated and detyrosinated microtubules is required for optimal growth, corresponding in our assay to 60%-70% of axon tips positively stained for detyrosinated tubulin (detyr^+^) (Fig. 1 A, B). Interestingly, parthenolide still reduced microtubule detyrosination and axon growth of VASH1 or VASH2 overexpressing neurons. However, to reach 60%-70% of detyr^+^ axon tips, at least ten times higher concentrations (10 nM) were required, while 1 nM parthenolide became insufficient to affect axon tip detyrosination and neurite growth of these neurons (Fig. 1 A-D). Moreover, concentrations ≥100 nM still slightly promoted axon growth in these cells (Fig. 1 C, D), indicating that higher parthenolide concentrations could counteract elevated microtubule detyrosination by VASH1 or VASH2 overexpression (Fig. 1 A, B). On the other hand, as GFP-controls, SVBP overexpressing neurons showed improved axon growth in the presence of 1 nM parthenolide (Fig. 1 C, D), suggesting that parthenolide directly targets VASH1 and VASH2.

### Endogenous VASH1/2 and SVBP mediate microtubule detyrosination in adult neurons

We then tested the contribution of endogenous VASH1, VASH2, and SVBP to the detyrosination of microtubules in axonal tips and generated baculoviruses for a knockdown in sensory neurons. Exogenously co-expressed HA- or FLAG-tagged proteins confirmed the shRNA constructs’ specificity and knockdown efficiency in sensory neurons (Fig. 1 E-J). Moreover, the efficient knockdown of endogenous VASH1, VASH2, or SVBP was verified by real-time PCR (Fig. 1 K-M) as specific antibodies for VASH1/2 or SVBP were unavailable for us.

After viral transduction of sensory neurons with these shRNA constructs, we replated them 5 days after culturing to ensure an efficient knockdown of each protein. Microtubule detyrosination and axon growth of replated cells were measured after an additional 24 h in culture. Interestingly, the knockdown of either VASH1, VASH2, or SVBP alone already significantly decreased the microtubule detyrosination in regenerated axons to about 50% of detyr^+^ axon tips (Fig. 1 N, O) and, therefore, beyond an optimal equilibrium of de- and tyrosinated microtubules for axon growth (similar to ineffective levels of 10 nM parthenolide) (Fig. 1 D). Consistently, additional parthenolide (1 nM) did not affect the axonal growth of these neurons (Fig. 1 P, Q). Thus, endogenous VASH1, VASH2, and SVBP contribute to microtubule detyrosination in regenerating axons. However, rather than pharmacological inhibition, their efficient knockdown already results in a too substantial reduction in microtubule detyrosination to promote axon regeneration.

To confirm this finding *in vivo*, we developed a method to transduce sensory and motor neurons with high efficiency in living mice. To this end, adult mice received a single intrathecal AAV1-GFP injection (Fig. 2 A), which transduced in average 92.5% of sensory neurons in L3-L5 DRG (Fig. 2 B-C). Moreover, AAV1-tdTomato injections, which resulted in similar transduction rates, enabled the detection of red-fluorescent axon tips in the footpad skin (Fig. 2 D). Additionally, 82.4% of neuromuscular junctions (NMJ) in the musculus extensor hallucis longus were innervated by labeled axons 2 weeks after injection (Fig. 2 E-F), indicating efficient transduction of motor and sensory neurons.

**Figure 2:**
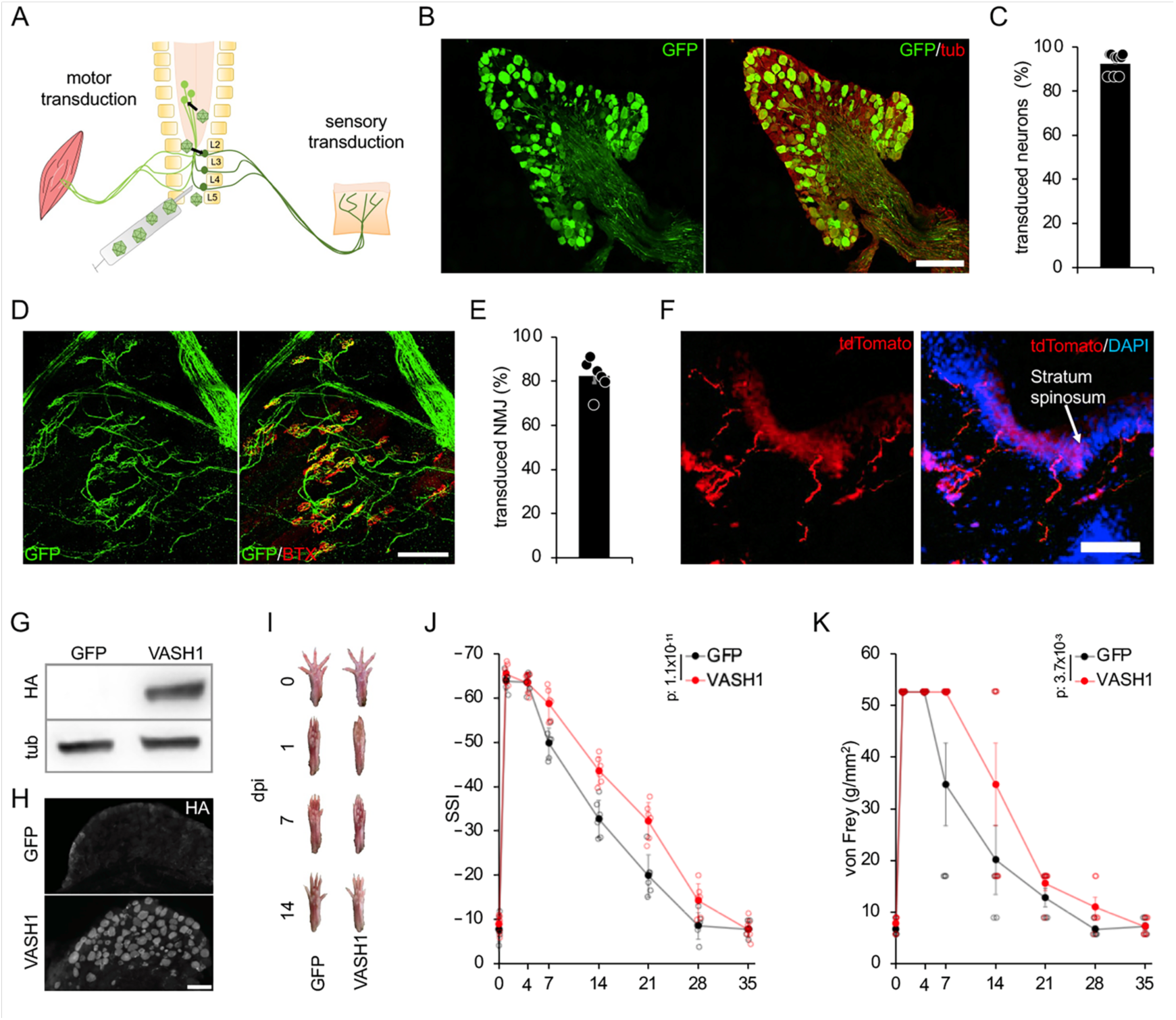
VASH1 overexpression reduces functional sciatic nerve regeneration. **A)** Schematic drawing depicting intrathecally injected AAV1 transducing motor neurons (left) and sensory neurons (right), whose axons project through the sciatic nerve to muscles or the footpad skin. **B)** L5 DRG section immunostained for GFP (green) and βIII-tubulin (tub, red). Scale bar: 200 µm. **C)** Quantification of transduced DRG neurons after intrathecal AAV1-GFP injection, showing transduction rates of >90%. The data represent mean ± SEM (bars) and single values (dots) of individual animals (n=9) animals **D)** Immunostaining of the muscle hallucis longus shows GFP-positive motor axons forming neuromuscular junctions (NMJ), identified by postsynaptic bungarotoxin-staining (BTX, red) 2 weeks after intrathecal AAV1 injection. Scale bar: 50 µm. **E)** Quantification of synapses innervated with GFP positive axons, 14 days after intrathecal AAV1-GFP injection, showing transduced innervation rates of 80%. The data represent mean ± SEM (bars) and single values (dots) of individual animals (n=7). **F)** Section of the footpads showing fibers from transduced sensory neurons (red) in the skin 2 weeks after intrathecal AAV1-tdTomato injection. The section was co-stained with DAPI. Scale bar: 50 µm. **G)** Western Blots of L3 and L4 DRG lysates of mice intrathecally transduced with either a GFP or a VASH1-HA containing AAV1. Western Blots were stained against HA and tubulin (tub) as a loading control. **H)** L4 DRG sections of mice intrathecally transduced either with GFP or VASH1-HA. Sections were immunostained for HA. Scale bar: 100 µm. **I)** Photos of right hind feet of mice before (0) or 1, 7, and 14 days (dpi) after sciatic nerve crush (SNC). Two weeks before SNC a VASH1 or GFP expressing AAV1 was injected intrathecally to transduce motor and sensory neurons projecting into the sciatic nerve **J)** Quantification of motor recovery in adult mice as depicted in I, using the SSI over 28 days after SNC. Each group contained six mice. Data represent the mean ± SEM. P-values were determined using two-way ANOVA followed by the Holm-Sidak post hoc test. **K)** Quantification of sensory recovery in the same mice described in I and J determined with the von Frey test. Data represent the mean ± SEM. P-values were determined using two-way ANOVA followed by the Holm-Sidak post hoc test.

We then used this approach for neuronal expression of either GFP (control) or VASH1 and performed a sciatic nerve crush (SNC) 2 weeks afterward (Fig. 2 G-K). Like in cultured sensory neurons, VASH1-expressing mice showed a significant delay in motor (assessed by the SSI) and sensory recovery after SNC (determined by the von Frey test compared to the corresponding non-diabetic controls) (Fig. 2 I-J). Overexpression of the HA-tagged VASH1 was verified via Western Blotting and immunohistochemistry (Fig. 2 G,H).

### Microtubule detyrosination and axon growth-promoting effect of parthenolide are age-dependent

As injured axons of young animals usually show a higher growth rate than adults (19–21), we investigated whether parthenolide can also accelerate axon growth further in postnatal neurons. To this end, we cultured sensory neurons from postnatal (P3) and adult (P70) mice. Postnatal neurons expectedly showed significantly longer axons than adult neurons after 48 h in culture (Fig. 3 A, B). However, parthenolide had no significant effect on axon growth in P3 cultures, while in P70 cultures, 1 nM of parthenolide increased axon growth to similar values as determined in postnatal cultures (Fig. 3 A, B). We, therefore, analyzed the degree of microtubule detyrosination in axon tips. Strikingly, axonal growth cones from postnatal neurons showed markedly lower microtubule detyrosination than adult cultures (P70) (Fig. 3 C, D), suggesting higher microtubule dynamics in growth cones of postnatal neurons. We then tested to what extent parthenolide treatment affected the detyrosination in these axons. While at 1 nM, parthenolide significantly reduced microtubule detyrosination in neurons from P70 mice to similar levels as in naïve postnatal neurons, it failed to decrease microtubule detyrosination in postnatal neurons further (Fig. 3 C, D). Thus, the higher detyrosination levels in adult neuron axonal growth cones compromise axonal growth. Moreover, a shift of microtubule detyrosination into the range of physiological levels of postnatal neurons facilitates axon growth of adult neurons.

**Figure 3:**
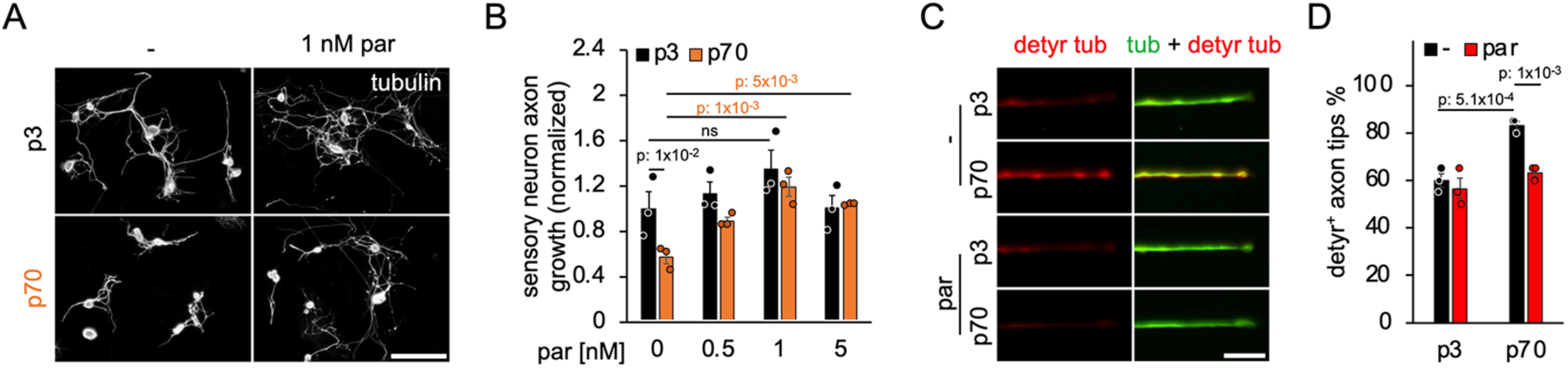
Postnatal neurons show reduced microtubule detyrosination in axon tips. **A)** Representative photos of axon tips of sensory neurons from postnatal (p3) and adult (p70) mice after 2 days in culture and treatment with either vehicle (−) or 1 nM parthenolide (par). Axons were stained for detyrosinated tubulin (detyr tub, red) and βIII-tubulin (tub). Scale bar, 5 μm. **B)** Quantification of detyrosinated tubulin in axon tips of cultured sensory neurons, as described in A. **C)** βIII-tubulin-stained neurons, as described in A, were treated with either 0, 0.5, 1, or 5 nM of par. Scale bar, 100 μm. **D)** Quantification of axon growth of neurons as depicted in C. Data from treated neurons were normalized to vehicle controls with an average axon length of 277 μm/neuron. Data represented in B and D show the mean ± SEM of three independent experiments. P-values were determined using two-way ANOVA followed by the Holm-Sidak post hoc test.

### Intravenous application of parthenolide and DMAPT promote nerve regeneration

A single injection of parthenolide into the sciatic nerve’s lesion site moderately enhances the length of regenerating axons in the distal segment when assessed 3 days after crush injury (11). To test whether a clinically more appropriate systemic application of parthenolide or one of its derivatives, DMAPT, is sufficient to facilitate axon growth in vivo, we applied either vehicle (PBS) or different doses of both drugs into adult mice’s tail vein daily, starting directly after sciatic nerve crush (SNC). Three days later, axon regeneration of superior cervical ganglion-10-positive (SCG10^+^) sensory neurons, choline-acetyltransferase-positive (CHAT^+^) motoneurons, and tyrosine hydroxylase-positive (TH^+^) sympathetic neurons were evaluated in immunostained longitudinal sciatic nerve sections (Fig. 4 A-F). While low doses of parthenolide (0.2 µg/kg) and DMAPT (2 µg/kg) showed only a slight effect on axon regeneration, results were more substantial at 2 µg/kg for parthenolide and 20 µg/kg for DMAPT, respectively (Fig. 4 A, B). At higher doses of 20 µg/kg the beneficial effect of parthenolide was decreased again (Fig. 4 A, B), confirming the bell-shaped dose-response curve previously reported in cell culture (11). Intravenous treatment with either parthenolide (2 µg/kg) or DMAPT (20 µg/kg) increased all neuronal subpopulations’ axonal lengths compared to respective vehicle-treated control groups (Fig. 4 A-F). TH+-axons were the longest among these different types of fibers. They reached up to 5 mm beyond the lesion site at this early time point, indicating higher axonal regeneration rates of sympathetic neurons compared to sensory and motor neurons.

**Figure 4:**
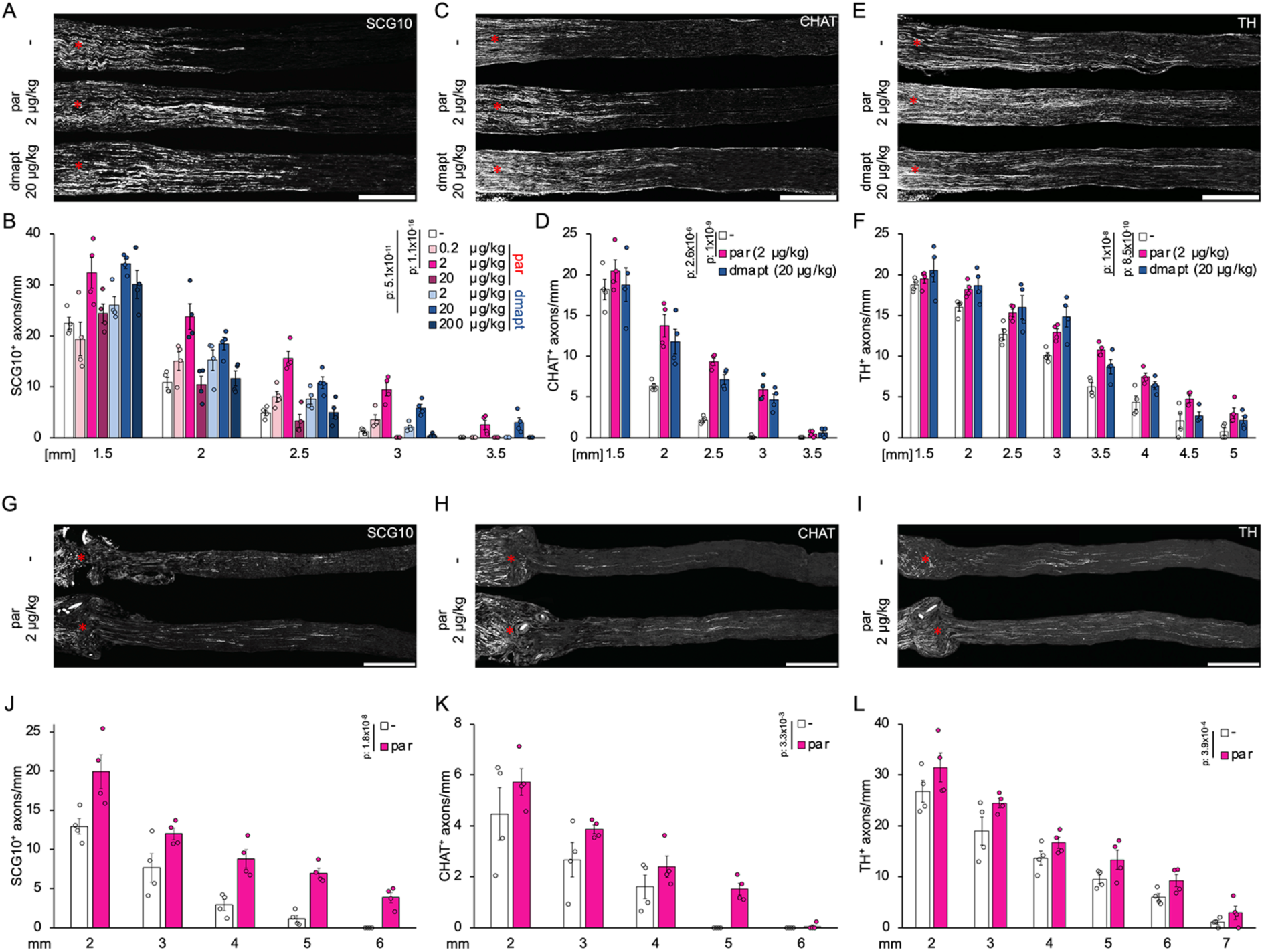
Systemic parthenolide or DMAPT applications accelerate the regeneration of different axonal types in various lesion models. **A)** Longitudinal sciatic nerve sections from adult mice stained with the sensory axon marker SCG10 3 days after crush injury. Animals had received daily injections of either vehicle (-), parthenolide (par), or DMAPT at indicated doses. Asterisks indicate the injury site. **B)** Quantification of regenerating axons at indicated distances beyond the injury site in the sciatic nerve of mice depicted in A. At least five sections per animal were analyzed from four animals per group. **C)** Longitudinal sections of the same animals described in A but stained for CHAT-positive motor axons with asterisks indicating the injury site. **D)** Quantification of regenerating axons at indicated distances beyond the injury site in the sciatic nerve of mice depicted in C. At least five sections per animal were analyzed from four animals per group. **E)** Longitudinal sections of the same animals described in A and C, stained for TH-positive sympathetic axons with asterisks indicating the injury site. **F)** Quantification of regenerating axons at indicated distances beyond the injury site of mice depicted in E. At least five sections per animal were analyzed from four animals per group. Scale bar for A, C, E, 500 μm. **G, H, I)** Longitudinal sciatic nerve sections from adult mice stained with either sensory (SCG10, G), motor (CHAT, H), or sympathetic (TH, I) axon markers, 7 days after nerve transection and anastomosis. Mice had been intravenously treated daily with vehicle (-) or parthenolide (par, 2 µg/kg). Asterisks indicate the injury site. Scale bar: 1 mm. **J, K, L)** Quantification of regenerating sensory (J), motor (K), and sympathetic (L) axons at indicated distances beyond the injury site in the sciatic nerves as depicted in G, H, and I, respectively. At least five sections per animal were analyzed from four animals per group. Data in B, D, F, J, K, L represent the mean ± SEM. P-values were determined using two-way ANOVA followed by the Holm-Sidak post hoc test.

As a systemic parthenolide application enabled axon growth promotion after a crush injury, we tested whether intravenously applied parthenolide enhances axon regeneration after complete nerve transection (neurotmesis) and anastomosis, where axonal regeneration is generally slower and less efficient than after nerve crush (3, 7, 22). To this end, we cut the sciatic nerve and performed surgical anastomosis. While controls, daily treated with vehicle, showed only a few short axons in the nerve’s distal part 7 days after injury, intravenously injected parthenolide (2 µg/kg/d) significantly increased the length of the sensory, motor, and sympathetic axons (Fig. 4 G-L). Thus, intravenous administration of parthenolide or DMAPT promotes the regeneration of sensory, motor, and sympathetic axons upon nerve injuries.

### Systemically applied parthenolide or DMAPT accelerate sensory and motor recovery in sciatic and median nerves

Next, we addressed whether intravenous application of parthenolide or DMAPT accelerates anatomic and clinically relevant functional recovery after nerve injury. To this end, we assessed motor and sensory function using the static sciatic index (SSI) and “von Frey” tests, respectively, over 4 weeks after SNC (8). Strikingly, daily repeated doses of parthenolide (2 µg/kg) or DMAPT (20 µg/kg) significantly improved the SSI score compared to vehicle-treated controls regarding both the onset of recovery and time to reach full recovery (Fig. 5 A-C). In contrast to vehicle-treated controls, parthenolide- and DMAPT-treated mice showed the first measurable improvement in motor function 4 days after injury (Fig. 5 A, B). In contrast, treatment effects concerning the touch response were first detectable 7 days after injury (Fig. 5 C). Consistent with the accelerated motor recovery by parthenolide, reestablished neuromuscular junctions (NMJ) were found in the *musculus extensor hallucis longus* (EHL) of parthenolide treated mice 4 days after SNC (Fig. 5 D). In contrast, very few reestablished NMJs were found in controls after this early time point. We also evaluated skin reinnervation in the footpads (Fig. 5 E-H). In cross-sections of the footpads, the axon numbers in the *stratum spinosum* (SS) dramatically dropped from an average of 28 per slice to 1 when assessed 1 day after SNC (Fig. 5 G, H), indicating extensive Wallerian degeneration. When evaluated 10 days after injury, parthenolide treatment had increased the number of axons in the SS layer significantly compared to the vehicle-treated control (Fig. 5 G, H). Thus, improved skin reinnervation was also correlated with parthenolide-mediated sensory recovery.

**Figure 5:**
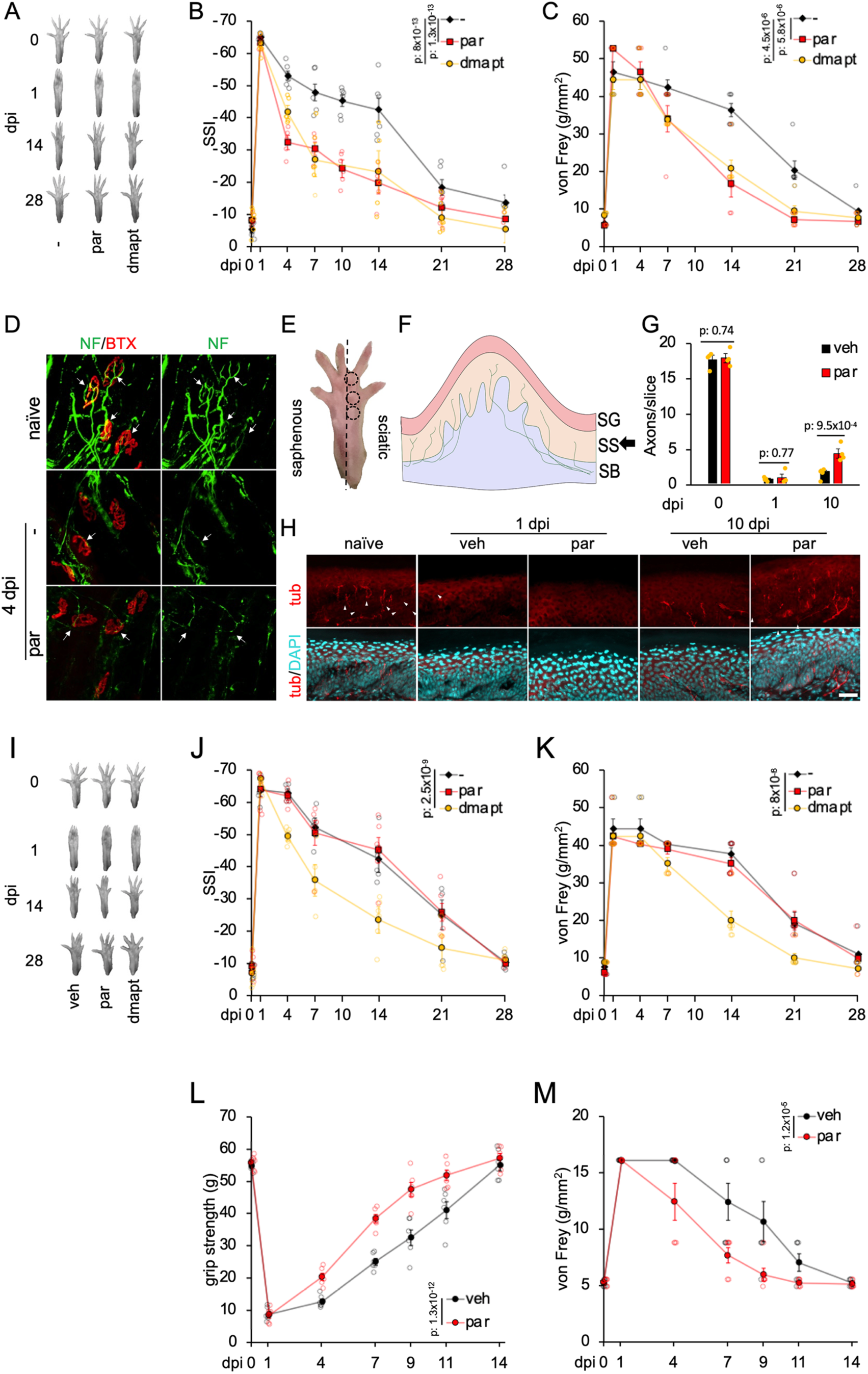
Systemic application of parthenolide or DMAPT accelerates target reinnervation and functional recovery. **A)** Photos of right hind feet of mice before (0) or 1, 14, and 28 days (dpi) after sciatic nerve crush (SNC). Animals had received daily intravenous doses of either vehicle (-), 2 µg/kg of parthenolide (par), or 20 µg/kg DMAPT. **B)** Quantification of motor recovery in adult mice as depicted in A, using the SSI over 28 days after SNC and daily intravenous injection with vehicle (-), parthenolide (par), or DMAPT. Each group contained six mice. Data represent the mean ± SEM. P-values were determined using two-way ANOVA followed by the Holm-Sidak post hoc test. **C)** Quantification of sensory recovery in the same mice described in A and B determined with the von Frey test. Data represent the mean ± SEM. P-values were determined using two-way ANOVA followed by the Holm-Sidak post hoc test. **D)** Representative pictures of *musculus extensor hallucis longus* (EHL) reinnervation after SNC + parthenolide (par) or vehicle (-) application. Muscular whole-mounts were stained for neurofilament (NF, green) and α-bungarotoxin (BTX, red). Arrows indicate reinnervated synapses. Scale bar, 50 µm. **E)** Areas of isolated footpads innervated by the sciatic (encircled) or saphenous nerves. **F)** Schematic drawing of a footpad cross-section showing the *stratum granulosum* (SG), *stratum spinosum* (SS), and *stratum basale* (SB). **H)** Footpad sections stained for βIII-tubulin (red) and DAPI (cyan). Arrowheads indicate axons in the SS. Scale bar, 10 µm. **G)** Quantification of the footpad reinnervation as depicted in H. Data represent mean ± SEM of four animals per group. Of each mouse, 25 slices per footpad were evaluated. P-values were determined using two-way ANOVA followed by the Holm-Sidak post hoc test. **I)** Representative photos of right hind paws of mice before (0) or 1, 14 and 28 days post-SNC (dpi). Mice orally received daily doses of a vehicle, 2 µg/kg par or 20 µg/kg DMAPT. **J)** Quantification of motor recovery in adult mice as depicted in A by the static sciatic index (n=6). Data represent the mean ± SEM. P-values were determined using two-way ANOVA followed by the Holm-Sidak post hoc test. **K)** Quantification of sensory recovery in adult mice determined by the von Frey test (n=6). Data represent the mean ± SEM. P-values were determined using two-way ANOVA followed by the Holm-Sidak post hoc test. **L)** Quantification of the grip strength of mice receiving either daily intravenous doses of vehicle (veh) or 2 µg/kg of parthenolide (par) over 14 days at the indicated time points before and after MNC. Each group contained six mice. **M)** Quantification of sensory recovery in adult mice described in L, determined with the von Frey test at the indicated time points before and after MNC. All data represent the mean ± SEM. All p-values were determined using two-way ANOVA followed by the Holm-Sidak post hoc test.

We then tested whether orally applied parthenolide or DMAPT is also effective. Daily applied oral DMAPT doses of 20 µg/kg fully mimicked the effects of intravenously injected DMAPT of similar dosages, while orally administered parthenolide showed no effect (Fig. 5 I-K). Even higher oral doses of parthenolide (20 µg/kg or 200 µg/kg) failed to improve nerve regeneration (suppl. Fig. 1 A, B), reflecting poor oral bioavailability of parthenolide (23).

Finally, we investigated whether the daily repeated intravenous applications of parthenolide (2 µg/kg) also promote other peripheral nerves’ regeneration. To this end, we crushed the right median nerve of mice and evaluated regeneration by the grip strength of the right front paw and the von Frey test. As for the injured sciatic nerve, parthenolide treatment significantly accelerated motor and sensory of the front paw (Fig. 5 L, M).

### Parthenolide accelerates functional nerve recovery in rats

After finding that parthenolide and DMAPT accelerate functional recovery in mice upon nerve injury, we investigated whether these drugs are effective in other species, where axons have to overcome longer regeneration distances to reach functional recovery. Therefore, we performed SNC in adult rats and treated them daily by intravenous applications of either vehicle or different parthenolide doses (0.2, 2, and 20 µg/kg/d), starting directly after SNC (Fig. 6 A-C). While daily doses of 0.2 and 20 µg/kg/d showed significantly but only moderate effects, efficacy was markedly higher at 2 µg/kg/d (similar to mice). The first effects could be seen in the SSI at this dosage 11 days after injury. In contrast, vehicle-treated rats showed first signs of recovery after 16 days (Fig. 6 B). Moreover, parthenolide-treated rats reached pre-injury SSI scores after 35 days, while the control group required another 14 days to reach these levels (Fig. 6 A, B). Sensory recovery was likewise accelerated upon intravenous parthenolide treatment (Fig. 6 C). Also, daily doses of 2 µg/kg showed the most potent effects: Improvements in the touch response were first detectable in this group after 14 days, while only seen in controls 21 days after injury. Furthermore, the function was restored to pre-surgery levels by parthenolide after 35 days, whereas the vehicle group took about 49 days (Fig. 6 C). In contrast to mice, SNC led to hypersensitivity in all rat groups, manifested earlier in parthenolide-treated animals. However, 56 days after SNC, vehicle-treated controls and groups treated with 0.2 or 20 µg/kg reached the same level of hypersensitivity (Fig. 6 C). Thus, optimal doses for intravenous parthenolide application in mice and rats were similar.

**Figure 6:**
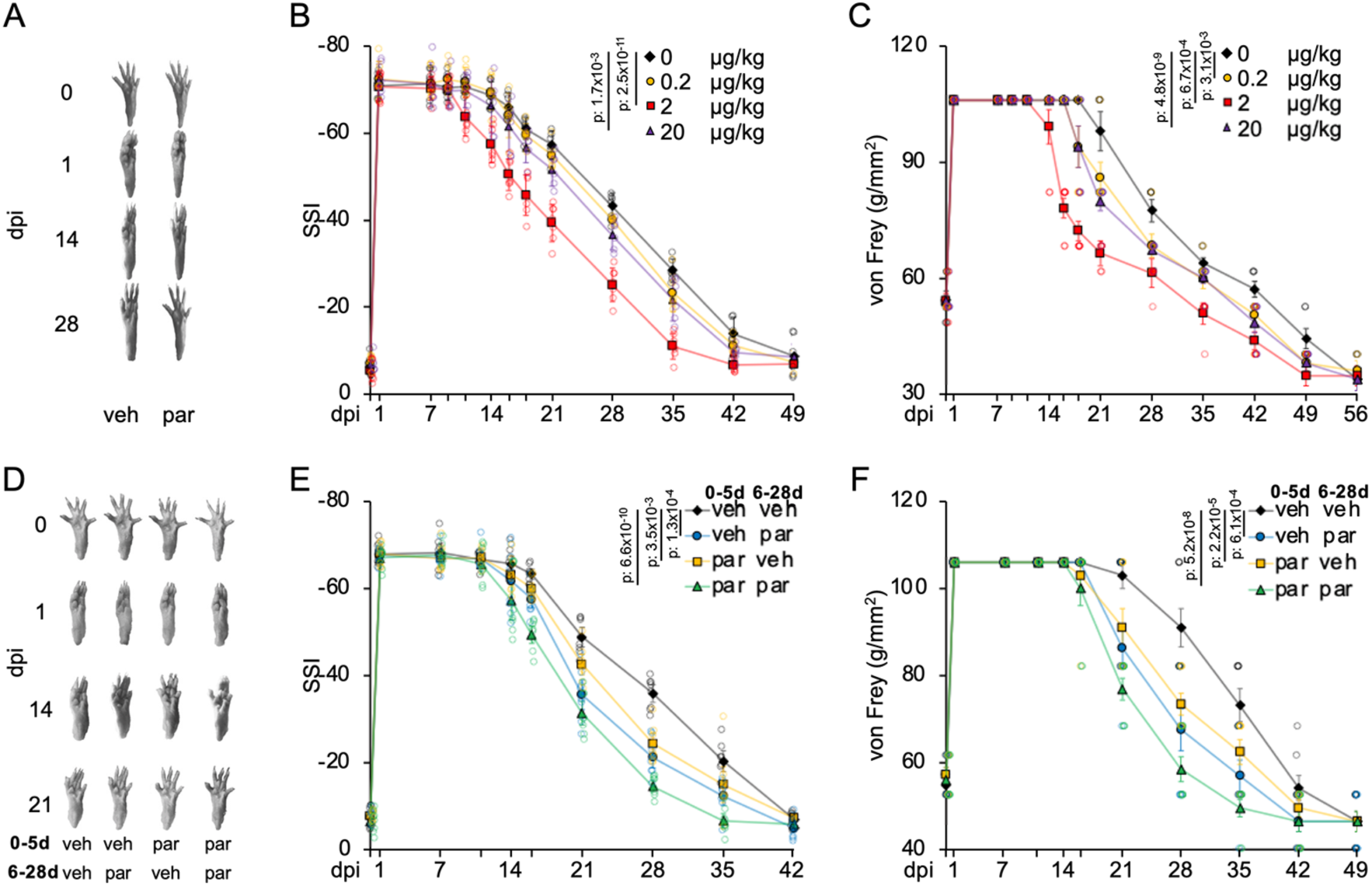
Systemic application of parthenolide promotes functional recovery in rats. **A)** Representative photos of right hind feet of rats before (0) or 1, 14, and 28 days after sciatic nerve crush (SNC; dpi). Adult rats received daily intravenous doses of either vehicle (0) or 2 µg/kg of parthenolide (par). **B)** Quantification of motor recovery in adult rats as depicted in A by the static sciatic index after SNC and daily intravenous injections of various doses 0.2 µg/kg, 2 µg/kg, or 20 µg/kg parthenolide over 49 days. Each group comprises six rats. Data represent the mean ± SEM. P-values were determined using two-way ANOVA followed by the Holm-Sidak post hoc test. **C)** Quantification of sensory recovery in the same rats described in B determined by the von Frey test. Data represent the mean ± SEM. P-values were determined using two-way ANOVA followed by the Holm-Sidak post hoc test. **D)** Representative photos of the right hind paw of rats before (0) or 1, 14, and 21 days after sciatic nerve crush (SNC; dpi) and different treatment regimes. Animals received daily treatment (intravenous injections) of either vehicle (veh) or parthenolide (par) from days 0-5 or 6-28 days after the injury as indicated. Another group received parthenolide treatment from day 0-28. **E)** Quantification of motor recovery in animals as depicted in D by the static sciatic index over 42 days after SNC. Each group contained six rats. Data represent the mean ± SEM. P-values were determined using two-way ANOVA followed by the Holm-Sidak post hoc test. **F)** Quantification of sensory recovery in the same rats described in D and E determined by the von Frey test. Data represent the mean ± SEM. P-values were determined using two-way ANOVA followed by the Holm-Sidak post hoc test.

### Delayed parthenolide treatment still improves functional recovery in rats

From a clinical point of view, it is relevant to know whether a delayed start of parthenolide treatment or just an initial treatment for some days is sufficient to accelerate functional recovery. As the onset of functional recovery occurs later in rats than in mice, we used adult rats and created four groups in which 2 µg/kg of parthenolide were intravenously administered daily using different treatment regimes. Animals of group 1 received the vehicle for 28 days; rats of group 2 received the vehicle for the first 5 days and then parthenolide for the remaining 23 days; animals of group 3 received parthenolide for the first 5 days and then the vehicle for the remaining 23 days; rats of group 4 received parthenolide daily for the entire 28 days. As expected, continuous treatment (group 4) showed the fastest motor and sensory recovery and group 1 (vehicle only) the slowest (Fig. 6 D-F). Moreover, animals of group 2 with a postponed treatment showed a delayed onset of functional improvement (Fig. 6 D-F) but only a slightly weaker effect at later time points than in group 4. Finally, animals of group 3, treated for the initial 5 days only, showed similar initial improvement as group 4 but were less effective than group 2 at later time points (Fig. 6 D-F). Thus, although continuous parthenolide treatment over the entire period showed the best results, a delayed start of the treatment was still effective.

## Discussion

Although peripheral nerves have, in principle, the ability to regenerate injured axons, functional recovery often fails to manifest due to a limited axonal growth rate. Thus, the clinical demand for drugs accelerating axon growth speed is high (7, 24). We have previously demonstrated that peripheral nerve regeneration in adult mice is accelerated in knockin mice with constitutively active GSK3 leading to inhibition of microtubule detyrosination (8, 11). Effects on microtubule detyrosination and axon growth in culture can be mimicked by the sesquiterpene lactone parthenolide (11). Here, we further established the relevant role of optimal levels of microtubule detyrosination to accelerate axon growth upon injury. While a substantial reduction of detyrosination in growth cones by parthenolide compromises axon growth, a relatively low decline enhances axon extension (Fig. 1 A-D, Fig. 3 A, B). However, the mechanism underlying the parthenolide effect on microtubule detyrosination and axon growth remained elusive. The enzymes vasohibin 1 (VASH1) and vasohibin 2 (VASH2) were recently identified as the first carboxypeptidases for microtubules (14–16, 25). Although parthenolide inhibits VASH activity (15, 16), a functional role in promoting axon growth remained elusive. Here we demonstrate that overexpression of VASH1 and VASH2, but not its enhancer SVBP increases microtubule detyrosination and compromises mature neurons’ axon regeneration. Moreover, the knockdown of endogenous VASH1, VASH2, or SVBP reduced detyrosination levels in axonal tips further than growth-promoting parthenolide concentrations (1 nM) and, therefore, parthenolide failed to accelerate axon growth in these neurons (Fig. 1 N, O). These data indicate the relevance of these enzymes on microtubule detyrosination in mature neurons and axonal growth. Furthermore, they illustrate that the axon growth-promoting effect of parthenolide relies on a relatively modest shift in the ratio of tyrosinated/detyrosinated microtubules, which lies in the physiological range of postnatal neurons. Accordingly, parthenolide failed to significantly accelerate the axon growth of postnatal neurons further (Fig. 3). On the other hand, parthenolide still promoted axon growth of VASH1, VASH2, and SVBP overexpressing neurons. However, at least 10-times higher drug concentrations were required to overcompensate for the higher VASH activity, while optimal concentrations for adult neurons (1 nM) were insufficient to affect axon growth in VASH overexpressing cells (Fig. 1 C, D). Based on these data and that parthenolide binds to VASH1 and VASH2 (16), we conclude that these proteins likely are the inhibited molecular targets mediating axon growth promotion. Although we cannot exclude that parthenolide also indirectly inhibits microtubule detyrosination by reducing the polymerization-competent pool of tubulin as reported recently (17), this possibility seems unlikely because the formation of adducts on cysteine and histidine residues on tubulin was reported at concentrations >1000-times higher than the concentrations used to promote axon regeneration (17). Furthermore, the low expression of VASH1 and VASH2 in neurons, the covalent binding of parthenolide to these proteins, and the relatively moderate shift of microtubule detyrosination required for accelerating axon growth are likely the reason for its high potency and its low dosages required to facilitate axon regeneration *in vivo*.

In addition, the current study’s finding that microtubule detyrosination is reduced in postnatal compared to adult sensory neurons and that reducing microtubule detyrosination in adult neurons to similar levels suggests that detyrosination is reduced may be a significant cause for the faster nerve regeneration observed in young compared to adult animals. So far, mainly axon extrinsic factors, such as diminished Schwann cell repair responses (20, 26), were considered responsible for the reduced regeneration in adult nerves. However, further research is required to explore whether age progression correlates with increased VASH or SVBP expression levels and understand why and how the degree of detyrosination is physiologically modulated in different neurons.

Currently, no therapeutics are available in the clinic to accelerate functional regeneration upon nerve injury. Therefore, our finding that intravenous injections or oral application of DMAPT accelerate functional recovery in different nerve injury paradigms and species is a promising step forward. In addition, the possibility for repeated applications over several weeks by these application routes is essential because, as shown in the current study, a continuous treatment was functionally more effective than a treatment over just a few days (Fig. 6 D-F). Furthermore, even a delayed onset of continuous parthenolide treatment was sufficient to accelerate functional recovery. As regeneration distances usually are longer in humans than in rodents, this aspect would be particularly relevant for potential clinical use in patients. Although it is currently unknown whether both drugs can also promote nerve regeneration in humans, their high potency and low effective dosages of 2 µg/kg (parthenolide) or 20 µg/kg (DMAPT) for promoting nerve regeneration in rats and mice make severe side effects unlikely, because DMAPT reportedly indicates no signs of toxicity or hematological parameters in mice when intravenously applied at 100 mg/kg (5,000-times higher than used in our study) over 10 consecutive days (23). Furthermore, since DMAPT accelerated functional sciatic nerve recovery at 5,000-fold lower doses (20 µg/kg), possible severe side effects at these dosages are most likely even more reduced. Similarly, administration of parthenolide at 25 mg/kg thrice weekly over 4 weeks or daily over 17 days showed no signs of toxicity (27, 28).

Oral drug administration is often preferable over intravenous treatment for routine everyday clinical therapy. Consistent with the low oral bioavailability of parthenolide, even much higher oral dosages of this drug remained ineffective. However, DMAPT, which reportedly has an oral bioavailability of about 80% (23), was effective at similar oral and intravenous dosages, making it particularly attractive as a medication. The higher required dosages of DMAPT compared to parthenolide are likely due to the covalent binding of parthenolide to a cysteine in the catalytic center of the VASH1/2 through C13. Thus, the thiol group of C169 reacts with the α-methylene of parthenolide through a Michael addition reaction (16, 29). As the C13 of DMAPT is reversibly bound to an amine, the Michael-addition must first dissociate back into parthenolide before becoming biologically active (30–32). Consistent with this, higher doses of the prodrug DMAPT were needed to accelerate axon growth *in vivo*.

In conclusion, the finding that systemic application of parthenolide or DMAPT accelerates functional recovery of injured nerves in different species opens the possibility for the first widely applicable medication to increase the critical time window for the reinnervation of peripheral targets. This means a faster return of motoric function and sensation and a better overall outcome after traumatic nerve injuries. Beyond that, these drugs may also help cure nerve diseases causing or associated with axonal damage, such as diabetic neuropathy (33, 34), axonal Guillain-Barre syndrome (35, 36), or chemotherapy-induced damage (34). Research investigating these possibilities is currently underway.

## Material and Methods

### Sensory neuron cultures and immunocytochemical staining

As described previously, sensory neurons were obtained from adult mice’s dissociated dorsal root ganglia (DRG) (8, 11). In short, isolated DRG (ca. T3-L6) were incubated in 0.25% trypsin/EDTA (GE Healthcare) and 0.3% collagenase Type IA (Sigma) in DMEM (Invitrogen) at 37°C and 5% CO2 for 45 min and mechanically dissociated. Cells were resuspended in DMEM containing 1:50 B27 (Thermo Fisher) and penicillin/streptomycin (500 U/ml; Merck, Millipore) and cultured at 37°C and 5% CO2 on poly-d-lysine (0.1 mg/ml, molecular weight <300 kDa; Sigma) and laminin (20 μg/ml; Sigma)-coated 8 well plates (Sarstedt). Cells were either treated with vehicle, 1-100 nM parthenolide. Axonal growth was determined after 48 h incubation by fixation in 4% PFA (Sigma) and immunocytochemical staining with antibodies against βIII-tubulin (1:2000; BioLegend, RRID:AB_2313773). Imaging was performed automatically with the Olympus VS210-S5 slide scanner. Total axon length and neuron numbers per well were automatically quantified with the NeuriteTracer plugin for ImageJ, avoiding experimenter-induced quantification bias. The average axon length per neuron and neuron counts per experimental group were normalized to control groups. Data represent the mean ± SEM of two to six independent experiments. Significances of intergroup differences were evaluated using two-way ANOVA followed by the Holm-Sidak post hoc test.

Microtubule detyrosination in axon tips was evaluated using antibodies against βIII-tubulin (1:2000; BioLegend, RRID:AB_2313773) and detyrosinated tubulin (1:500; Sigma, RRID:mAB_477583) as described before (37, 38). Axon tips were defined as the last 15 μm of βIII-tubulin-positive neurite extensions. Data represent mean ± SEM of three replicated wells with 30–60 tips per well from three independent experiments. Significances of intergroup differences were evaluated using two-way ANOVA followed by the Holm-Sidak *post hoc* test.

### Surgical procedures

All animal protocols adhered to animal care guidelines and were approved by the local authorities (LANUV Recklinghausen). Male and female adult (8-12 weeks) C57/BL6J mice and Lewis rats were maintained in cages with 1-5 animals on a 12 h light/dark cycle with ad libitum access to food and water. Sciatic nerve crush (SNC) was performed as described previously (8, 11). In brief, mice were anesthetized by intraperitoneal injections of ketamine (80-100 mg/kg, Pfizer) and xylazine (10-15 mg/kg, Bayer), rats with 2% isoflurane in oxygen, and a skin incision of ∼10 mm was made above the gluteal region. Then, the ischiocrural musculature was carefully spread with minimal tissue damage to expose the right sciatic nerve from the sciatic notch to the point of trifurcation. Next, the crush injury was performed for 10 s proximal to the tibial and peroneal divisions using graphite powder-dipped Dumont #5 forceps (Hermle) to mark the crush site. The skin was then closed using 6-0 sutures.

A skin incision of ∼10 mm from the right axillary region to the elbow was made for the mouse median nerve crush, thus exposing the median nerve from its origin at the brachial plexus to the elbow. Subsequently, the crush injury was performed for 10 s using graphite powder-dipped Dumont #5 forceps (Hermle) to mark the crush site. The skin was then closed using 6-0 sutures. After the surgery, animals received daily intravenous or oral doses of parthenolide or DMAPT in 100 µl vehicle.

### Intrathecal AAV1 injection

For intrathecal AAV application, a 5 mm skin incision was made above the L4-L5 spines. The muscle tissue was carefully spread, and 2.5 µl of the viral solution was injected into the cerebrospinal fluid between the L4 and L5 spines using a Nanoject III injector (Drummond Scientific) with three consecutive 833 nl injections with a speed of 7 nl/s and 20 s intervals between injections. The skin was closed using 6-0 sutures. Transduction was performed 2 weeks before sciatic nerve crush (SNC).

### Quantification of regenerating axons in the sciatic nerve

Sciatic nerves were isolated 3 d after SNC, postfixed in 4% PFA overnight, dehydrated in 30% sucrose at 4°C again overnight, and embedded in Tissue-Tek (Sakura). Longitudinal sections (14 μm) were cut on a cryostat (Leica), thaw-mounted onto coated glass slides (Superfrost Plus, Fisher), and stored at −20°C. Cryosections were immunohistochemically stained with antibodies against the regeneration-associated sensory axon marker SCG10 (1:1000; Novus Biologicals, RRID:AB_10011569) (39), the motor axon marker CHAT (1:100; Sigma, RRID:AB_90661) or the sympathetic axon marker Tyrosine Hydroxylase (1:500; Novus Biologicals, RRID:AB_10077691). In addition, labeled axons were quantified beyond the graphite-labeled injury site, as previously described (8, 11). Experimental groups comprised four to five animals, and five different sections were analyzed per animal. Statistical significances of intergroup differences were evaluated using two- or three-way ANOVA followed by the Holm–Sidak post hoc test.

### Immunohistochemical DRG staining

Mouse L4 DRG were removed 2-6 weeks after intrathecal AAV1 transduction and fixed in 4% PFA over night at 4°C. Followlingly, DRG were dehydrated in 30% sucrose at 4°C again overnight, and embedded in Tissue-Tek (Sakura). Sections (14 μm) were cut on a cryostat (Leica), thaw-mounted onto coated glass slides (Superfrost Plus, Fisher), and stored at −20°C. Cryosections were immunohistochemically stained with antibodies against βIII-tubulin (1:2000; BioLegend, RRID:AB_2313773), GFP (1:500, Novus, RRID:AB_10128178), or HA (1:500, Sigma-Aldrich, RRID:AB_2600700) overnight at 4°C.

### WESTERN BLOT ASSAYS

L3 and L4 DRG from three mice per experimental group were combined in 100 μl lysis buffer (20 mm Tris-HCl, pH 7.5, 10 mm KCl, 250 mm sucrose, 10 mm NaF, 1 mm DTT, 0.1 mm Na3VO4, 1% Triton X-100, 0.1% SDS, and protease inhibitors; Merck, Millipore) and homogenized by sonification. Lysates were centrifuged at 8000 × g for 10 min at 4°C to remove cell debries, and soluble proteins were separated by SDS-PAGE. Subsequently, they were transferred to nitrocellulose membranes (Bio-Rad) according to standard protocols. Blots were blocked either in 5% dried milk (Sigma) and incubated either with monoclonal antibodies against βIII-tubulin (1:5000; BioLegend, RRID:AB_2313773) or HA (1:500, Sigma-Aldrich, RRID:AB_2600700) overnight at 4°C. Antibodies were detected with anti-rabbit or anti-mouse IgG secondary antibodies conjugated to HRP (1:80,000; Sigma). Antigen-antibody complexes were visualized using enhanced chemiluminescence substrate (Bio-Rad) on the Fluorchem E imaging system (ProteinSimple).

### Baculoviral vectors and knockdown

Plasmids: HA-tags were attached to mouse VASH1 (Origene, MR222520) and VASH2 (Origene, MR203958) by PCR (vash1-HA fw: cgatcgccatgccaggggg; VASH1-HA rev: atccgggtgtacccatacgatgttccagattacgcttaactcgagaaaa; VASH2-HA fw: cgatcgccatgtggctgcacg; VASH2-HA rev: gatccggatctacccatacgatgttccagattacgcttaactcgagaaaa. FLAG-tagged SVBP was cloned from cDNA derived from murine heart tissue by PCR. FLAG-SVBP fw: gactacaaagacgatgacgacaagatggatccacctgcc; FLAG-SVBP rev: gcagccgcctggggagtgactcgag. For SVBP or VASH1/2 knockdown primers containing the shRNA sequence SVBP-shRNA fw: gatccgatgagttctgtaagcagatgctcgagcatctgcttacagaactcatcttttta; SVBP shRNA rev: agcttaaaaagatgagttctgtaagcagatgctcgagcatctgcttacagaactcatcg; VASH1sh fw: gatccgctgtgatcctgggaatttacctcgaggtaaattcccaggatcacagctttttaa; VASH1sh rev: agctttaaaaagctgtgatcctgggaatttacctcgaggtaaattcccaggatcacagcg; VASH2sh fw: gatccgacttcgaagattcctataagctcgagcttataggaatcttcgaagtcttttta; VASH2sh rev: agcttaaaaagacttcgaagattcctataagctcgagcttataggaatcttcgaagtc) were cloned into U6 plasmids.

Baculovirus production was performed as described recently (18, 40): According to the manufacturer’s protocols, recombinant baculoviruses were produced using the ViraPower BacMam Expression System (Thermo Fisher). In brief, recombinant bacmid DNA was purified and transfected into adherent Sf9 cells (Thermo Fisher) using Cellfectin reagent to generate P1 recombinant baculovirus stock. Baculoviruses were amplified by inoculation of 50 ml Sf9 suspension cultures (10^6^ cells/ml) in Sf-900 III SFM Medium supplemented with 12.5 U/ml penicillin/streptomycin (Biochrom) in 125 ml polycarbonate Erlenmeyer flasks with vent cap (Corning) with 1 ml virus stock solution and incubation at 27°C and 110 rpm for 4 days. Baculovirus preparations were pre-tested on HEK293 cells (seeded at ∼3–5 × 10^4^ cells per well in 96-well plates) by adding 10 μl virus per well overnight. Transduction efficiencies of ≥ 90% were regarded as appropriate for further use. Otherwise, virus stock was subjected to further amplification cycles. Knockdown efficiencies were verified by immunostaining against HA (1:500; Sigma-Aldrich, RRID:AB_2600700) for VASH1 and VASH2 or FLAG (1:500; Sigma, RRID:AB_262044) for SVBP after baculoviral transduction in sensory neurons. The staining intensity was calculated using the formula intensity=integrated density - area * mean gray value background. Transduction was verified by immunostaining against GFP (1:500; Novus; RRID:AB_10128178), and neurons were identified by βIII-Tubulin staining (1:1000; BioLegend; RRID:AB_2313773). Additionally, the knockdown was verified by quantitative real-time PCR (qPCR) with cDNA from sensory neurons after baculoviral induced VASH1/2 or SVBP knockdown after 5 days in culture. For neuronal purification, sensory neurons were cultivated in the presence of 1 µM arabinofuranosylcytosine and 20 µM fluor-deoxyuridine to eliminate dividing cells. qPCRs primers as followed: VASH1 for: TGGCCAAGATCCACCCAGATG; VASH1 rev: TCGTCGGCTGGAAAGTAGGCAC; VASH2 for: AGGGGGAGAGATGGTAGGCGC; VASH2 rev: AGCCAGTCTGGGATCGTCATGG; SVBP for: AACCAGCCTTCAGAGTGGAGAAGG; SVBP rev: GCTCCGTCATGACTCTGTTGAGAGC.

### Adeno-associated Virus

AAV1-GFP was obtained from Addgene (#37825-AAV1) with a 7×1012 vg/ml titer. The other AAV1 viruses were produced in our laboratory using a pAAV-IRES_eGFP expression vector for VASH1-HA. AAV plasmids carrying either cDNA for respective genes downstream of a CMV promoter were co-transfected with pAAV2/1 (Addgene #112862) encoding the AAV genes rep and cap, and the helper plasmid (Stratagene) encoding E24, E4, and VA into AAV-293 cells (Stratagene) for recombinant AAV1 generation. Virus particles were purified as described previously (41). The titer of the AAVs ranged from 1.2 × 10^14^-2 × 10^14^ GC/ml. Mainly neurons are transduced upon intrathecal injection of AAV1, as this virus serotype is highly neurotropic.

### Static sciatic index (SSI)

The static sciatic index was determined as described recently (8, 42, 43): After SNC, motor recovery was determined by calculating the SSI, as previously described (8). At the same time of day (11:00 a.m. - 2:00 p.m.) at various time points after SNC (mice: 0, 1, 4, 7, 9, 14, 21 and 28 d; rats: 0, 1, 7, 9, 11, 14, 16, 18, 21, 24, 28, 30, 32, 35, 37, 42, 45, 49, 53 and 63 d), mice or rats were lifted from the ground to photograph the left and right hind feet, respectively. The treatment was blinded to the experimenter during the experiment. Toe spreading on the contralateral (C, left) and the ipsilateral (I, right) sides relative to the SNC was assessed by measuring paw length (PL) and the distance between the first and fifth toe (FF). The SSI was then calculated using the previously described formula: SSI = 101.3 ((IFF − CFF)/CFF) − 54.03((IPL - CPL)/CPL) - 9.5 (43). Data represent mean ± SEM per experimental group. Statistical significances of intergroup differences were evaluated using two-way ANOVA followed by the Holm-Sidak post hoc test.

### Grip strength

After the median nerve crush (MNC), the right forepaw’s functional motor recovery was assessed by determining the grip strength return. Therefore, the grip strength was tested at various time points after MNC (0, 1, 4, 7, 9, 11, and 14 d) at the same time of day (11:00 a.m. - 2:00 p.m.).

For the test, the mouse was held by the tail close enough to the bar of the computerized grip strength meter (Model 47230, Ugo Basile, Varese, Italy) to grasp it. When the mouse held onto the bar, it was carefully pulled away until losing grip. The metal bar was connected to a force transducer that automatically recorded each measurement’s peak force in grams. At each time point, five measurements were recorded. Of the five measurements, the mean of the three middle values was determined as the grip strength. Statistical significances of intergroup differences were evaluated using two-way ANOVA followed by the Holm-Sidak post hoc test.

### Von Frey test

The von Frey test was determined as described recently (8, 11, 44):

Sensory recovery after SNC and MNC was determined at various time points (mice, SNC: 0, 1, 4, 7, 9, 14, 21, 28 d; mice, MNC: 0, 1, 4, 7, 9, 11 and 14 d; rats, SNC: 0, 1, 7, 9, 11, 14, 16, 18, 21, 24, 28, 30, 32, 35, 37, 42, 45, 49, 53 and 56 d) after SNC with the von Frey filament test as previously described. Tests were performed at the same time of day by the same experimenter, unaware of the treatment. Mice or rats were placed on an elevated metal grid (pore size: 2 mm) and allowed to acclimate for 15 min before testing. Starting with the smallest, differently sized, innocuous von Frey filaments (Muromachi Kikai) were consecutively poked into the ipsilateral hind paw to elicit a paw withdrawal. Statistical significances of intergroup differences were evaluated using two-way ANOVA followed by the Holm-Sidak post hoc test.

### Analysis of muscle reinnervation after SNC

Mice were sacrificed before crush or 4 days after SNC. The extensor hallucis longus muscle was dissected, fixed in 4% PFA for one hour, and permeabilized in 2% Triton-X/PBS overnight. Axons were labeled with an antibody against heavy and medium-chain neurofilament (1:2000; Cellsignaling) and Alexa488-conjugated secondary antibody. Reestablished synapses in neuromuscular endplates were visualized by incubation with Alexa594-conjugated α-BTX (1:1000; Invitrogen) for 1 h at room temperature. Representative pictures of each experimental group were taken on an SP8 confocal microscope (Leica).

### Analysis of footpad reinnervation after SNC

The sciatic nerve of mice was crushed. Mice were injected intravenously with 2 µg/kg parthenolide (Sigma) or vehicle (DMSO) daily 10 days before tissue isolation. The footpads innervated by the sciatic nerve were isolated. The tissue was harvested from uninjured animals and animals 1 and 10 days after sciatic nerve crush. The tissue was fixed in 4% PFA at 4°C overnight and dehydrated in 30% sucrose at 4° C again overnight. 20 µm cryosections were permeabilized in methanol and blocked (2% BSA, 5% donkey serum). Axons were identified by immunostaining against βIII-tubulin (1:1000, Covance) overnight, and antibodies were detected with anti-rabbit antibodies conjugated with Alexa-594. In addition, nuclei were stained with DAPI (Merck).

Axons in the *stratum spinosum* of 25 sections per mouse were counted. Four mice per condition were analyzed. The experimental conditions were blinded for the experimenter. Statistical significances of intergroup differences were evaluated using two-way ANOVA followed by the Holm-Sidak post hoc test.

## Acknowledgments

We thank Anastasia Andreadaki, Nadine Hube, Swenja Henne, Christopher Brennsohn, Kessy Brzozowski, Laura-Marie Twadorwski, and Celina Böse for technical support. The Federal Ministery of Education and Research supported this work.

---

**Supplementary Fig. 1:**
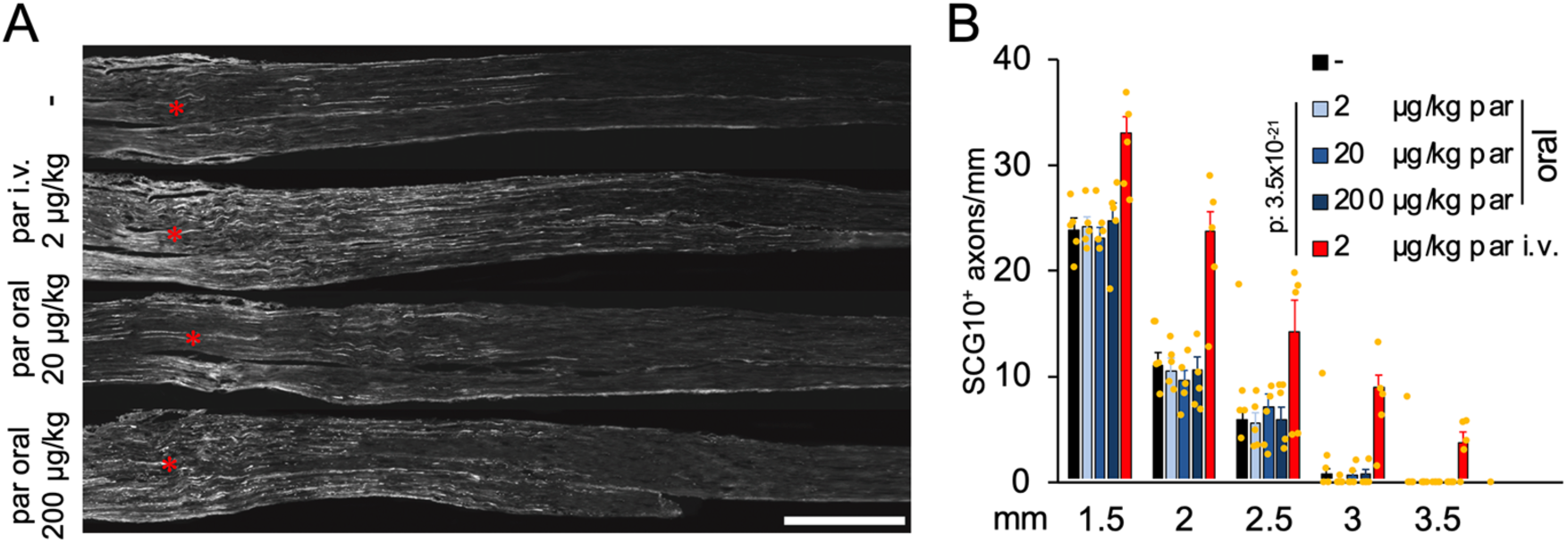
Orally applied parthenolide fails to increase anatomic sciatic nerve regeneration. **A)** Longitudinal sciatic nerve sections from adult mice stained with the sensory axon marker SCG10 3 days after crush injury. Animals had received daily injections of either vehicle (-) or parthenolide (par) at indicated doses. Asterisks indicate the injury site. **B)** Quantification of regenerating axons at indicated distances beyond the injury site in the sciatic nerve of mice depicted in A. At least five sections per animal were analyzed from five animals per group. Data represent the mean ± SEM. P-values were determined using two-way ANOVA followed by the Holm-Sidak post hoc test.

